# Potent macrocycle inhibitors of the human SAGA deubiquitinating module

**DOI:** 10.1101/2021.05.13.444038

**Authors:** Michael Morgan, Tatsuya Ikenoue, Hiroaki Suga, Cynthia Wolberger

**Affiliations:** Department of Biophysics and Biophysical Chemistry, Johns Hopkins University School of Medicine, Baltimore, MD 21205 USA; Graduate School of Science, The University of Tokyo, Hongo, Bunkyo-ku, Tokyo 113- 0033, Japan

**Author notes:** Corresponding author: Cynthia Wolberger, Hiroaki Suga.

## Abstract

The SAGA transcriptional coactivator contains a four-protein subcomplex called the DUB module that removes ubiquitin from histone H2B-K120. The human DUB module contains the catalytic subunit, USP22, which is overexpressed in a number of cancers that are resistant to available therapies. We screened a massive combinatorial library of cyclic peptides and identified potent inhibitors of USP22. The top hit was highly specific for USP22 as compared to a panel of 44 other human DUBs. Cells treated with peptide had increased levels of H2B monoubiquitination, demonstrating the ability of the cyclic peptides to enter human cells and inhibit H2B deubiquitination. These macrocycle inhibitors are, to our knowledge, the first reported inhibitors of USP22/SAGA DUB module and show promise for development.

## Introduction

Histone ubiquitination serves a non-degradative role in regulating transcription (Zhang, 2003, Henry et al., 2003, Turner et al., 2002), DNA repair (Moyal et al., 2011), DNA replication (Somasagara et al., 2017, Hung et al., 2017), and chromatin condensation (Fierz et al., 2011). Monoubiquitination of histone H2B (H2B-Ub) at lysine 120 in humans and lysine 123 (Robzyk et al., 2000) in yeast is a hallmark of actively transcribed genes (Weake and Workman, 2008). H2B-Ub plays a variety of roles in transcription, including promoting assembly of the pre-initiation complex (PIC) (Kao et al., 2004, Turner et al., 2002) and stimulating nucleosome reassembly by the histone chaperone, FACT (Fleming et al., 2008, Pavri et al., 2006). H2B-Ub also stimulates H3K4 methylation by human MLL1 and yeast COMPASS (Sun and Allis, 2002, Dover et al., 2002, Xue et al., 2019, Kim et al., 2009), and H3K79 methylation by human Dot1L and yeast Dot1 (Briggs et al., 2002, McGinty et al., 2008, Steger et al., 2008, Ng et al., 2002a, Ng et al., 2002b). H2B-Ub is distributed throughout gene bodies and is particularly enriched at transcription start sites (TSSs), (Kao et al., 2004, Schulze et al., 2011). An intriguing aspect of H2B-Ub is that it is a transient mark that is rapidly turned over during transcription, with both ubiquitination and deubiquitination of H2B important for maintaining wild type levels of transcription (Batta et al., 2011, Henry et al., 2003, Kao et al., 2004). Deubiquitination of histone H2B plays a role in promoting phosphorylation of the RNA polymerase II CTD by Ctk1, thereby promoting the elongation phase of transcription (Wyce et al., 2007). Studies have shown that genes with low H2B-Ub levels are expressed at higher levels than those with high H2B-Ub (Prenzel et al., 2011, Hahn et al., 2012, Shema et al., 2008). Consistent with findings in yeast, surveys of gene expression in primary human cancer cells found that loss of USP22 resulted in changes in transcription at SAGA regulated genes (Atanassov et al., 2016).

Altered levels of the enzymes that both attach and remove H2B-Ub are implicated in oncogenesis. Ubiquitin is covalently linked to H2B-K120 in human cells by the E2 ubiquitin conjugating enzyme, hRAD6A/B (Koken et al., 1991) and the E3 ubiquitin ligase, RNF20/40 (Kim et al., 2005). Misregulation of both the E2 and E3 enzymes that monoubiquitinate H2B have been shown to result in overexpression of genes that drive cell proliferation (Hung et al., 2017, Shema et al., 2008, Somasagara et al., 2017). A number of different deubiquitinating enzymes (DUBs) remove ubiquitin from H2B at distinct subsets of genes, including USP22, USP51, USP27x, USP36, and USP44 (Fuchs et al., 2012, Atanassov et al., 2016, DeVine et al., 2018). Of these DUBs, altered expression of USP22 has been implicated in a variety of tumor types (Melo-Cardenas et al., 2016). USP22 overexpression has been identified as part of an 11- gene “death by cancer” signature of gene expression – a pattern observed over diverse tissue types that is characteristic of metastatic cancers that respond poorly to existing therapies (Glinsky et al., 2005). USP22 overexpression correlates with poor clinical outcomes in tumors of the brain (Li et al., 2013), breast (Zhang et al., 2011), stomach (Yang et al., 2011), liver (Liao et al., 2019, Wen et al., 2020), and colon (Liu et al., 2010, Liu et al., 2012), although its mechanistic role in oncogenesis is poorly understood. In addition, reduced expression of USP22 is associated with chromosomal instability (CIN), since efficient chromosome compaction during mitosis requires USP22 deubiquitination of H2B in metaphase (Jeusset et al., 2021). Silencing of USP22 expression results in aberrant chromosomal segregation and phenotypes consistent with CIN, including polyploid daughter cells (Jeusset et al., 2021). Taken together, these findings suggest that appropriate USP22 activity is a critical regulator of major cellular events, and its misregulation is correlated with oncogenesis and genomic heterogeneity.

Because of its role in cancer, USP22 is an attractive target for drug discovery. USP22 is the enzymatic DUB subunit of the SAGA transcriptional coactivator complex, a 1.8 MDa complex comprising 19 subunits that are organized into functional modules (Zhang et al., 2008, Soffers and Workman, 2020). The SAGA complex is conserved from yeast to humans and binds to promoter regions, where SAGA facilitates assembly of the transcriptional pre-initiation complex (Grant et al., 1998a, Grant et al., 1998b, Massari et al., 1999, Sterner et al., 1999), acetylates histone H3 and deubiquitinates histone H2B (reviewed in (Soffers and Workman, 2020)). The deubiquitinating activity of SAGA resides in a four-protein subcomplex called the DUB module (Lee et al., 2009, Kohler et al., 2006, Kohler et al., 2008, Lang et al., 2011, Ingvarsdottir et al., 2005). As first shown in yeast, the catalytic subunit alone is inactive unless incorporated into a complex containing the other three DUB module subunits (Rodriguez-Navarro et al., 2004, Ingvarsdottir et al., 2005, Kohler et al., 2008, Ellisdon et al., 2010). The yeast DUB module comprises the catalytic subunit, Ubp8, in complex with Sgf73, Sgf11, and Sus1 (Henry et al., 2003, Ingvarsdottir et al., 2005, Rodriguez-Navarro et al., 2004, Kohler et al., 2010, Samara et al., 2010). Crystal structures of the yeast DUB module revealed a remarkably intertwined subunit arrangement in which each subunit contacts the other three (Kohler et al., 2010, Samara et al., 2010). Deletion or disruption of the interface between Ubp8 and the zinc finger domain of Sgf11, which anchors the DUB module to the nucleosome acidic patch (Morgan et al., 2016), destabilizes intersubunit interactions and abrogates enzymatic activity (Morgan et al., 2016, Samara et al., 2012, Yan and Wolberger, 2015, Wang et al., 2020, Kohler et al., 2010). The human DUB module contains the catalytic subunit, USP22, in complex with ATXN7 (homologue of Sgf73), ATXN7L3 (homologue of Sgf11), and ENY2 (homologue of Sus1) (Zhao et al., 2008, Zhang et al., 2008). Like its yeast homologue, USP22 depends on its partner subunits for full activity (Atanassov et al., 2016), which makes it necessary to utilize the full human DUB module in screens for novel inhibitors.

We report here the results of a screen for macrocyclic peptide inhibitors that target USP22. We took advantage of the RaPID (**Ra**ndom non-standard **P**eptides **I**ntegrated **D**iscovery) system, which comprises a massive combinatorial library of structurally unique cyclic peptides that can be screened in a high-throughput manner based on binding affinity. Using the human SAGA DUB module containing USP22 bound to its three partner proteins, we identified six cyclic peptides that bind tightly to the DUB module and inhibit DUB module activity at sub-micromolar concentrations on both fluorogenic and nucleosomal substrates. The most potent peptide inhibitor, which has an apparent Ki of about 20 nM on the human DUB module, did not inhibit 45 other human DUBs, indicating that it is highly specific for USP22. Remarkably, this inhibitor also showed greater activity on USP22 than on two other DUBs, USP27x and USP51, that also deubiquitinate histone H2B and form complexes with two of the SAGA DUB module adaptor subunits. Cells treated with the cyclic peptide inhibitors had increased H2B-Ub levels, indicating that the peptides can also inhibit USP22 in vivo. These compounds are promising leads as treatments for cancers in which USP22 overexpression may be a factor.

## RESULTS

### Screen for macrocyclic peptides that inhibit the DUB module

We employed the RaPID system to screen libraries of over 10^12^ unique macrocyclic peptides (Hipolito and Suga, 2012, Ito et al., 2015) to identify candidate inhibitors based on their ability to bind tightly to the human DUB module (hDUBm) or yeast DUB modules (yDUBm) (Figure 1A). A puromycin-ligated mRNA library was constructed to encode peptides with *N*-chloroacetyl-L-Tyrosine (^L^Y-library) or *N*-chloroacetyl-D- Tyrosine (^D^Y-library) as the initiator amino acid, followed by a random peptide region consisting of 6–15 residues, a cysteine and ending with a short linker peptide. Upon translation of these mRNAs, the chloroacetyl group on N-terminus of the linear peptides spontaneously cyclizes with the downstream cysteine to form thioether-macrocyclic peptides. The cyclic scaffold ensured that the ^L^Y-library and the ^D^Y-library diversified three-dimensional structures. Each cyclic peptide was covalently linked to its corresponding mRNA template via the puromycin linker for later amplification and DNA sequencing. These libraries were applied to streptavidin-conjugated magnetic beads to which a N-terminally biotinylated hDUBm or yDUBm was immobilized for selection of ligands against each enzyme complex.

**Figure 1.**
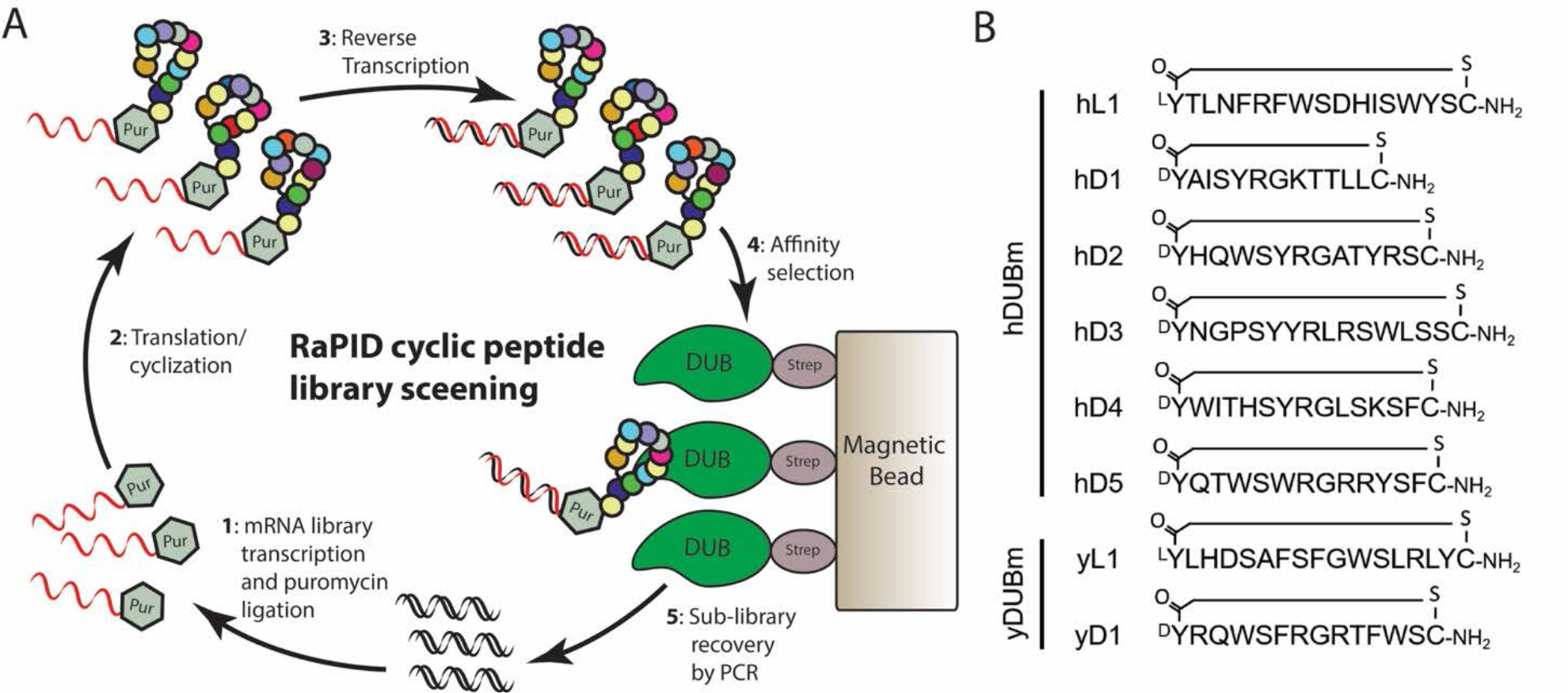
RaPID cyclic peptide library screening: **A)** Schematic of Random nonstandard Peptide Integration Discovery platform used for targeting the SAGA DUB module for novel binding species. **B)** Sequences for each peptide selected from RaPID screen for further evaluation as inhibitors. Each peptide contains the indicated D- or L-amino acid at the N- terminus covalently linked to the C-terminal cysteine sulfhydryl group.

Following four or five rounds of selection, the recovery rate of peptide–mRNA fusion molecules was significantly increased (Figure S1), suggesting that the population of cyclic peptides binding hDUBm or yDUBm were selectively enriched. Sequence analysis of the respective enriched libraries yielded unique sequences for the peptides. We selected the six most enriched cyclic peptides for hDUBm and two sequences targeting yDUBm on the basis of the hit frequency for further analysis.

### Peptides are potent inhibitors of the human DUB module

The cyclic peptides that constituted potential inhibitors were chemically synthesized by solid phase peptide synthesis and tested for their effectiveness and specificity as inhibitors in vitro. We assayed hDUBm activity using the fluorogenic substrate, ubiquitin- amino methyl coumarin (Ub-AMC) over a range of inhibitor concentrations. As shown in Figures 2A–2F, all six peptides that were identified based on tight binding to the human DUB module were also potent inhibitors of DUB activity. All peptides had sub- micromolar Ki values, with the hL1, hD1, hD2, and hD3 peptides exhibiting values of < 100 nM. The most potent inhibitor as judged by RaPID screening and *K*i approximation (Kuzmic et al., 2000), hD1, had an apparent Ki of 20 nM.

**Figure 2.**
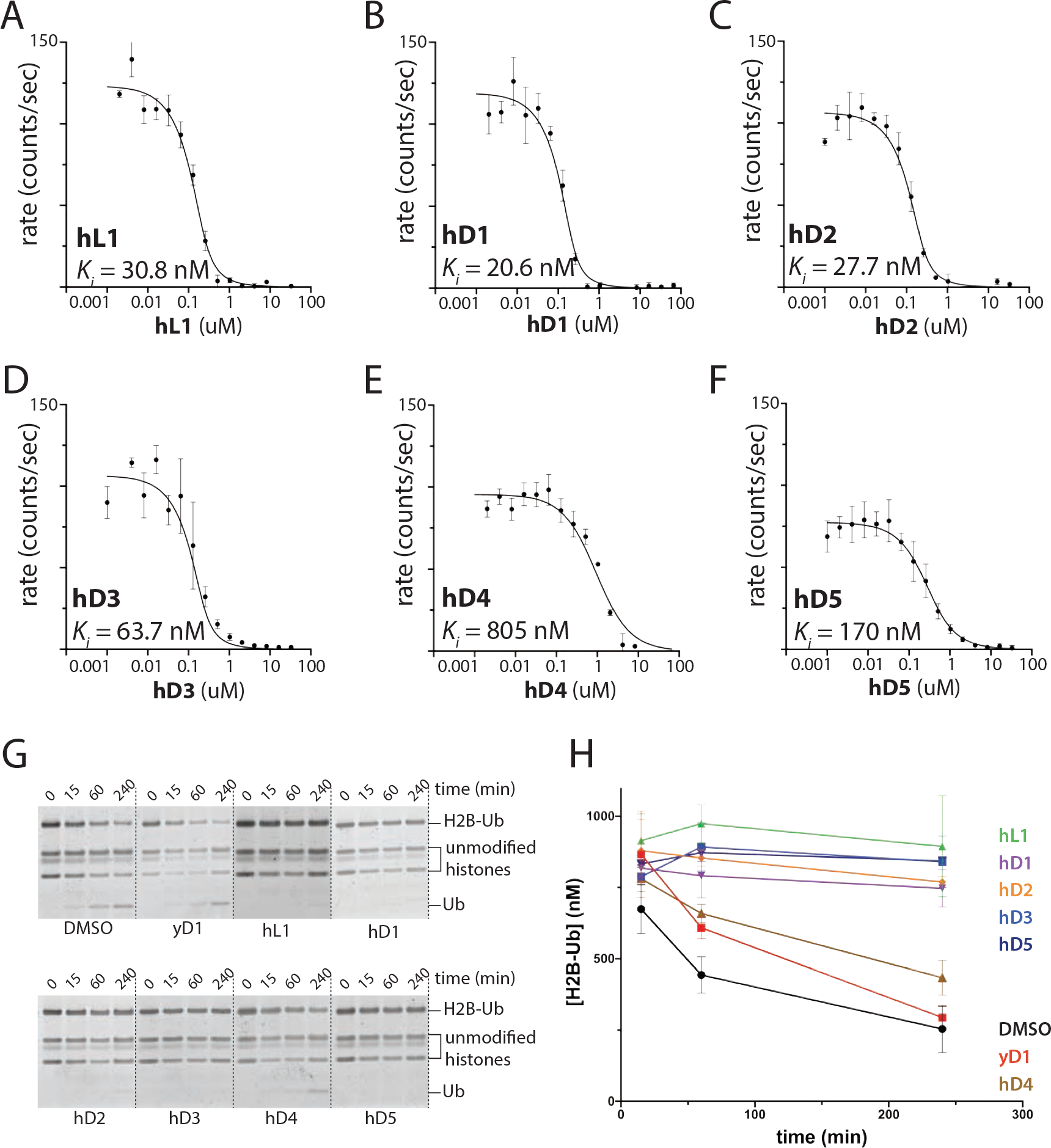
Inhibition of human DUB module activity inhibition by cyclic peptides. **A-F)** Each indicated cyclic peptide was tested for its ability to inhibit Ub-AMC cleavage in the presence of increasing amounts of the peptide into a reaction with a fixed amount of DUB (200 nM) and Ub- AMC (1 µM). *K*i values were fitted using the Morrison approximation with a fixed *K*m value of 35 µM as determined in Figure 3A. **G)** Inhibition of human DUB module activity on H2B ubiquitinated nucleosomes. Effect of each cyclic peptide (1 uM) and DMSO control on hDUB module cleavage rates on ubiquitinated nucleosomes. **H)** Quantitation of the disappearance of the H2B-Ub band, performed in triplicate.

The two peptides targeting the yeast DUB module were not pursued as inhibitors, as yL1 was insoluble in aqueous conditions and yD1 did not inhibit activity of the yeast DUB module (Figure S2A). Since yD1 also did not inhibit the human DUBm (Figure S2B), nor was it enriched in the screen for peptides that bind tightly to the human DUBm, this peptide was used for control experiments.

To test the effectiveness of the inhibitors on a native-like substrate, we assayed the ability of each cyclic peptide to inhibit DUB activity on nucleosomes ubiquitinated at H2B-K120. A denaturing gel was used to monitor cleavage of H2B-Ub, releasing H2B and Ub (Figure 2G). With the exception of hD4, all of the cyclic peptide leads were potent inhibitors of nucleosomal H2B-Ub cleavage by the human DUB module (Figures 2G and 2H). Each cyclic peptide sufficiently inhibited hDUB module activity such that no substantial activity was detected over the time course examined. The peptide with the highest Ki on Ub-AMC, hD4 (805 nM) (Figure 2E), also showed very weak inhibition of hDUBm on nucleosomal H2B-Ub. These results confirm the ability of cyclic peptides hL1, hD1, hD2, hD3, and hD5 to inhibit the enzymatic activity of USP22/DUB module on a native substrate.

### Characterization of inhibition mechanism

There are a variety of possible mechanisms by which the macrocycles could potentially inhibit the USP22/DUB module. We analyzed the inhibition kinetics of the most potent inhibitor, hD1, to determine the mode of inhibition (Fig. 3A). DUBm activity as a function of substrate concentration was assayed in the presence of several different concentrations of hD1 and then fit the data to competitive, non-competitive, and uncompetitive models of inhibition. We obtained the best fit with a non-competitive model, suggesting that hD1 does not compete for ubiquitin binding, but instead exerts its effect on the catalytic step of the reaction. Data from these experiments indicate that the Ki of hD1 is ∼180 nM, while Km was calculated to be 35 µM and kcat was 2.8 s^-1^, matching previous experiments.

**Figure 3.**
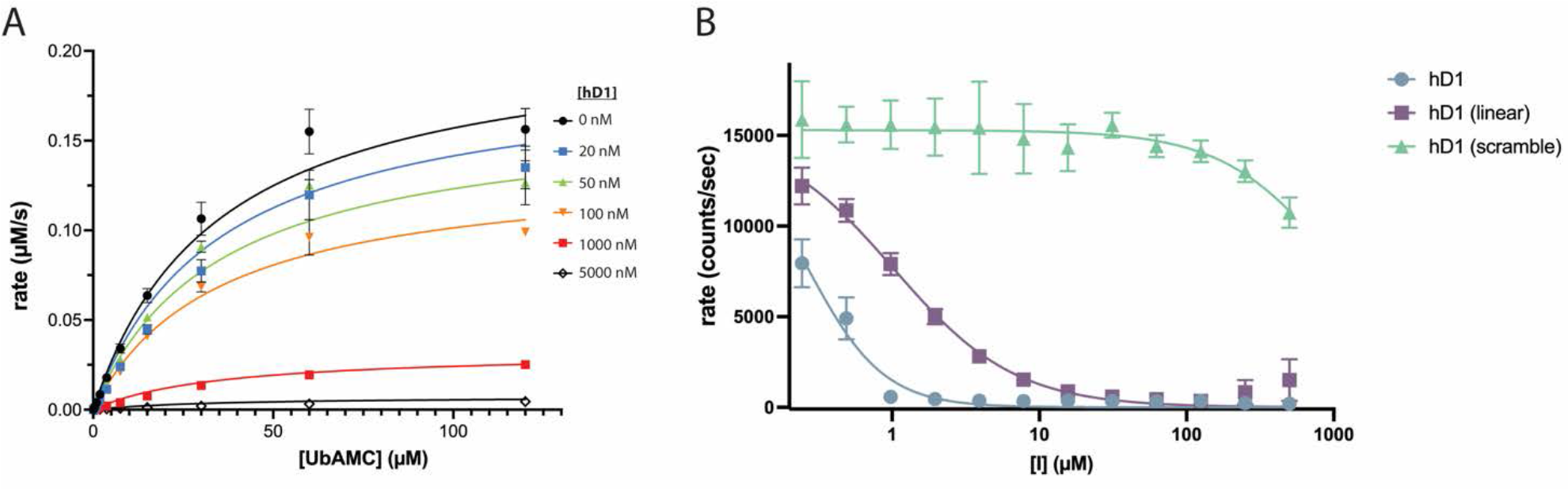
hD1 is a non-competitive inhibitor requiring the cyclized form for optimal USP22 targeting. **A)** hDUBm cleavage rates were measured at multiple substrate and hD1 concentrations to determine inhibitor equilibrium binding constant (*K*i) and mode of inhibition. The data were best fit to a non-competitive model of enzyme inhibition, with a measured *K*m of 35 µM, kcat of 2.8 s^-1^, and *K*i of 180 nM. **B)** Cleavage rates of 5 µM Ub- AMC by 200 nM hDUBm in the presence of the indicated concentration of either linear, scrambled, or cyclic hD1 peptide sequences fitted using IC50 analysis. IC50 values were calculated as 270 nM for the cyclic hD1, 1 µM for the linear version, and >1000 µM for the scrambled hD1 sequence.

A rationale for using small, cyclized peptides as inhibitors is that cyclization constrains the conformation of the peptide backbone, thus reducing the unfavorable entropic penalty upon formation of a complex with the target enzyme. To test the importance of peptide cyclization, we tested the ability of a linear hD1 peptide to inhibit USP22. As shown in figure 3B, linear hD1 inhibits USP22 at a ∼4-fold higher concentration, indicating the constraining the peptide does not play a major role in the effectiveness of this inhibitor. To verify that the peptide sequence itself is important, rather than overall amino acid composition, we tested a peptide of the same length but with a scrambled amino acid sequence and showed that it failed to inhibit DUB activity, even at very high peptide concentrations (Fig. 3B).

Most inhibitors disrupt activity by binding directly to the catalytic domain either in or near the enzyme active site. Since USP22 is only active when bound to the other DUB module proteins (Lang et al., 2011, Atanassov et al., 2016), another possible mechanism by which the cyclic peptides inhibit USP22 activity could be by altering the interactions between USP22 and the three other subunits, ATXN7, ATXN7L3, and ENY2. Studies of the yeast DUB module showed that deletion of the Sgf11 zinc-finger leads to an inactive domain-swapped dimer of DUB modules (Samara et al., 2012), while mutations made at the interface between Ubp8 and the Sgf11 zinc finger significantly reduce enzyme activity (Kohler et al., 2010). Similarly, disruption of intersubunit interactions between the catalytic subunit, Ubp8 (yeast homologue of USP22), and Sgf73 (homologue of ATXN7) cause a loss of DUB activity (Yan and Wolberger, 2015). To see if the USP22 inhibitors altered the overall organization of human DUB module subunits, we used small-angle x-ray scattering (SAXS) to determine whether the DUB module undergoes a large-scale change to its shape in the presence each respective cyclic peptides. Since there is no structure of the human DUB module, we compared the results to predicted SAXS data calculated from the crystal structure of the yeast DUB module (PDB ID: 3MHH) (Samara et al., 2010).

SAXS data were measured with the human DUB module incubated with each respective peptide inhibitor, using DMSO as a vehicle control. The Kratky plot (Fig. 4A) and *p(r)* distribution (Figure 4B) of the human DUB module in the presence of the hD1 inhibitor are very similar to those of the yeast complex, consistent with the idea that the two complexes adopt a similar tertiary and quaternary structure. A similar trend is seen in SAXS data collected on the human DUB module in the presence of the other cyclic peptides (Figure S3), suggesting that the peptides appear to bind the DUB module at discrete surfaces, rather than disrupting the complex. Consistent with this conclusion, ab initio density maps calculated for the human and yeast DUB modules in the presence and absence of hD1 using DENSS analysis (Grant, 2018) indicated minor changes in the molecular envelope. Interestingly, binding of each cyclic peptide slightly increased the long axis of the complex (Figure 4C). The greatest difference in molecular envelope due to peptide binding occurs in the region of the Sgf11/ATXN7L3 zinc finger, which is located at one end of the long axis (Figure 4C). We conclude from these data that all the peptide-bound human DUB module complexes largely maintain their fold at peptide concentrations that inhibit enzymatic activity.

**Figure 4.**
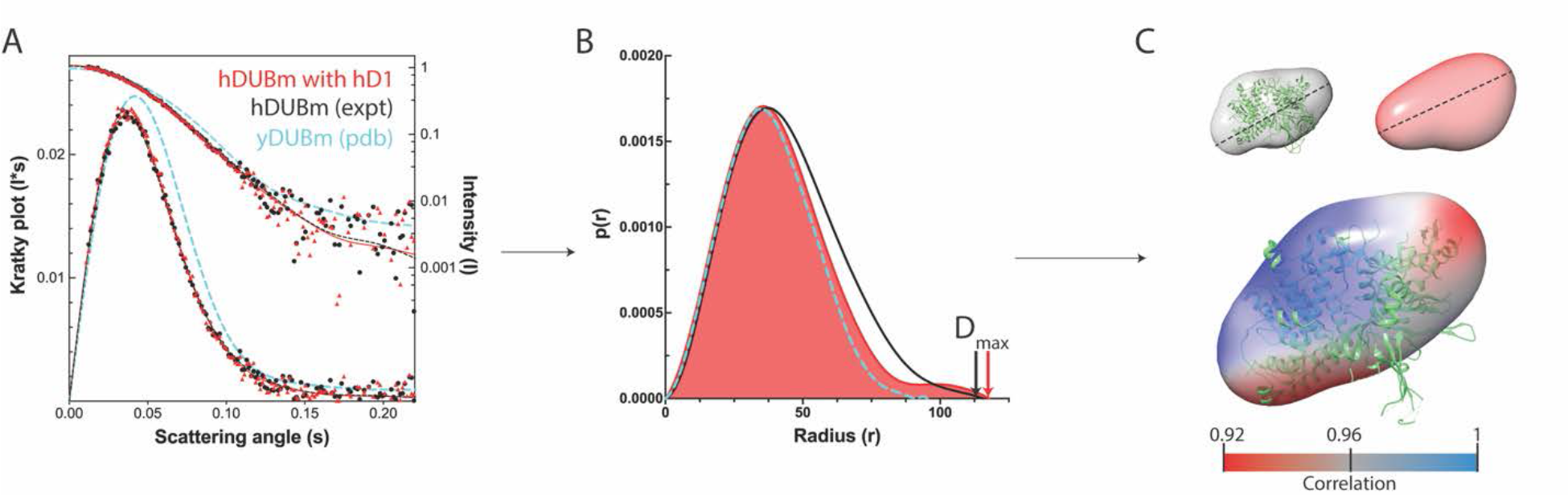
Small angle x-ray scattering (SAXS) analysis of the DUB module in the presence and absence of cyclic peptide. **A)** Scattering intensities of 1 mg/mL (6 µM) hDUBm alone (black), hDUBm in the presence of 7uM hD1 (red), and theoretical scattering intensities calculated from the yDUBm crystal structure (PDB ID: 3MHH, cyan). The right y-axis describes raw measured intensities (upper curve), while the left y-axis is a plot of I*s values (Kratky plot). **B)** Frequency distribution (P(r)) of possible vectors between surfaces on the scattering sample. Experimental curves from hDUBm (black) and hDUBm plus hD1 inhibitor (red); calculated curve for yDUBm (cyan). D_max_ is indicated by a colored arrow corresponding to each sample. **C)** Electron density maps derived from SAXS data of hDUBm alone (gray) and hDUBm in the presence of hD1 (red). Dashed line indicates D_max_. Correlations between the peptide-bound and apo complexes indicated by pseudocoloring (scale below). The yeast DUB module structure (green) was used to independently orient both density maps.

### hD1 is highly selective for USP22

The human genome encodes about 80 active DUBs (Clague et al., 2015), 55 of which belong to the same USP structural family as USP22, thus presenting a challenge to the development of specific DUB inhibitors. To probe the relative specificity of hD1, we screened a panel of 44 human DUBs for inhibition, including 27 USP DUBs (DUB*profiler*^TM^; Ubiquigent). The activity of each DUB was tested in the presence of a single concentration of inhibitor and compared with enzyme activity in the absence of compound. We chose 1 µM as the concentration of hD1 to test, as the USP22/DUB module has no detectable activity under these conditions (Figure 2B). As shown in Figure 5A, hD1 is strikingly specific: it did not significantly affect the activity of any of the enzymes tested except for USP27x, whose activity was reduced by ∼50%. Interestingly, USP27x has been reported to deubiquitinate histone H2B-K120 in vivo and in vitro as part of a complex with ATXN7L3 and ENY2 (Atanassov et al., 2016), two of the three adapter proteins that are part of the USP22 DUB module. In this screen, however, USP27x alone was tested in the absence of other subunits.

**Figure 5.**
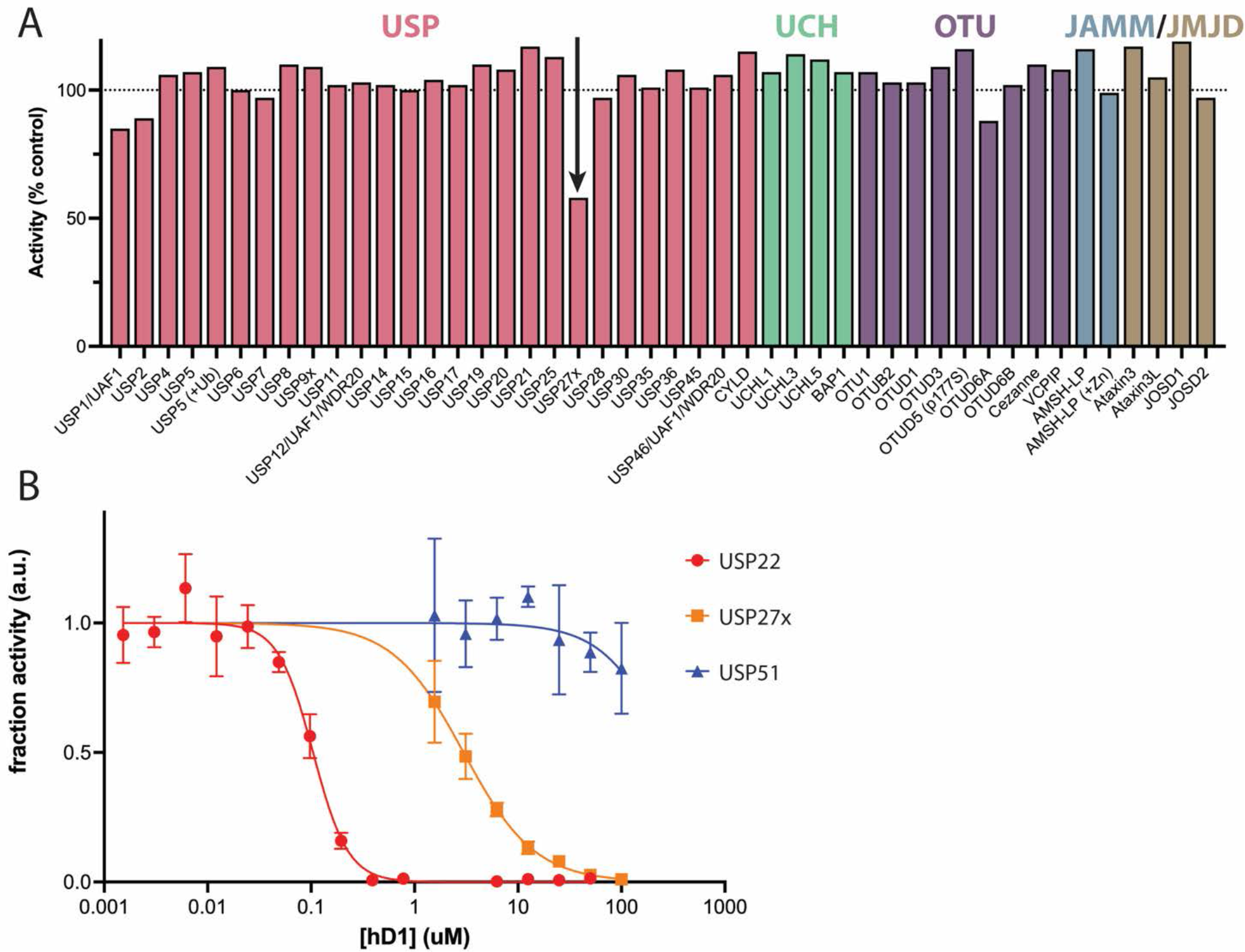
hD1 is specific for USP22. **A)** DUBProfiler assay of the effect of hD1 on a panel of 44 human DUBs. Cleavage of a ubiquitin-rhodamine(110)-glycine substrate was compared in the presence and absence of 1 µM hD1. **B)** Inhibition of H2B DUB complexes as a function of hD1 concentration. Cleavage rates of 5 µM Ub-AMC were assayed for 75 nM USP22/ATXN7L3/ENY2/ATXN7 (hDUBm), 100 nM USP27x/ATXN7L3/ENY2, or 1000 nM USP51/ATXN7L3/ENY2. Rates were normalized to the uninhibited reaction in each case to determine the fraction of apparent activity. IC50 values of hD1 were 100 nM for USP22, 3 µM for USP27x, and >250 µM for USP51.

We next compared the relative specificity of hD1 on USP22, USP27x, and USP51 in the context of their native complexes. We purified heterotrimeric complexes containing USP27x and USP51, respectively, in complex with ENY2 and ATXN7L3. The activity of these complexes on Ub-AMC was compared to that of the human DUBm, which contains USP22, ENY2, ATXN7L3, and a fourth subunit, ATXN7, that anchors the DUBm to the rest of the SAGA complex (Kohler et al., 2008, Herbst et al., 2021). As shown in Figure 5B, hD1 inhibits the USP27x complex at >25-fold higher inhibitor concentrations and modestly inhibits the USP51 complex only at >2500-fold higher concentration than that needed to inhibit the human DUB module. The high specificity of hD1 for USP22 as compared to USP27x and USP51 is all the more notable given that all three complexes deubiquitinate nucleosomal histone H2B-K120Ub. In addition, the finding that hD1 inhibits USP27x alone as well as the USP27x/ENY2/ATXN7L3 complex suggest that hD1 inhibits DUB activity by binding directly to the catalytic domain, rather than to the ENY2 and ATXN7L3 subunits. We note that an analogous comparison with USP22 and USP51 was not feasible because the two proteins are poorly behaved in solution when expressed on their own.

### Cyclic peptide inhibitor increases H2B ubiquitination in cells

We next asked whether each inhibitor could increase levels of H2B ubiquitination in cells, which would be consistent with inhibition of USP22. We introduced each cyclic peptide as a supplement to the growth medium of HEK293T cells, extracted the histones from each culture after two hours of exposure, and quantitated levels of H2B- Ub by immunoblot. Levels of H2B-Ub were compared to control cells that were treated with DMSO, the yD1 cyclic peptide that has no effect on USP22 activity, or the broad-spectrum cell permeable DUB inhibitor, PR-619. The relative abundance of histone H3 was used to control for differences in nuclear extraction. As shown in Figure 6A, treatment with five of the six inhibitors significantly increased H2B-Ub levels as compared to the controls. The most dramatic results were observed for hD1 and hD3, which increased H2B-Ub levels in excess of 15-fold, followed by hD4 and hD5, which increased H2B-Ub levels by about 12-fold (Figure 6B). The hD2 and hL1 peptides resulted in a minor increase of H2B-Ub on the scale seen with the yD1 control peptide and the DUB inhibitor, PR-619. The robust effect of hD4 in the cell-based assay is interesting given that this was the least effective inhibitor in the Ub-AMC cleavage assay, with an apparent Ki of ∼800 nM. Overall, the accumulation of H2B-Ub indicates that the inhibitors can enter cells and inhibit DUB module activity. While we cannot rule out an effect on other DUBs that are specific for H2B-Ub, such as USP27x and USP51 (Atanassov et al., 2016), our results are consistent with the ability of these inhibitors to inhibit USP22 in cells.

**Figure 6.**
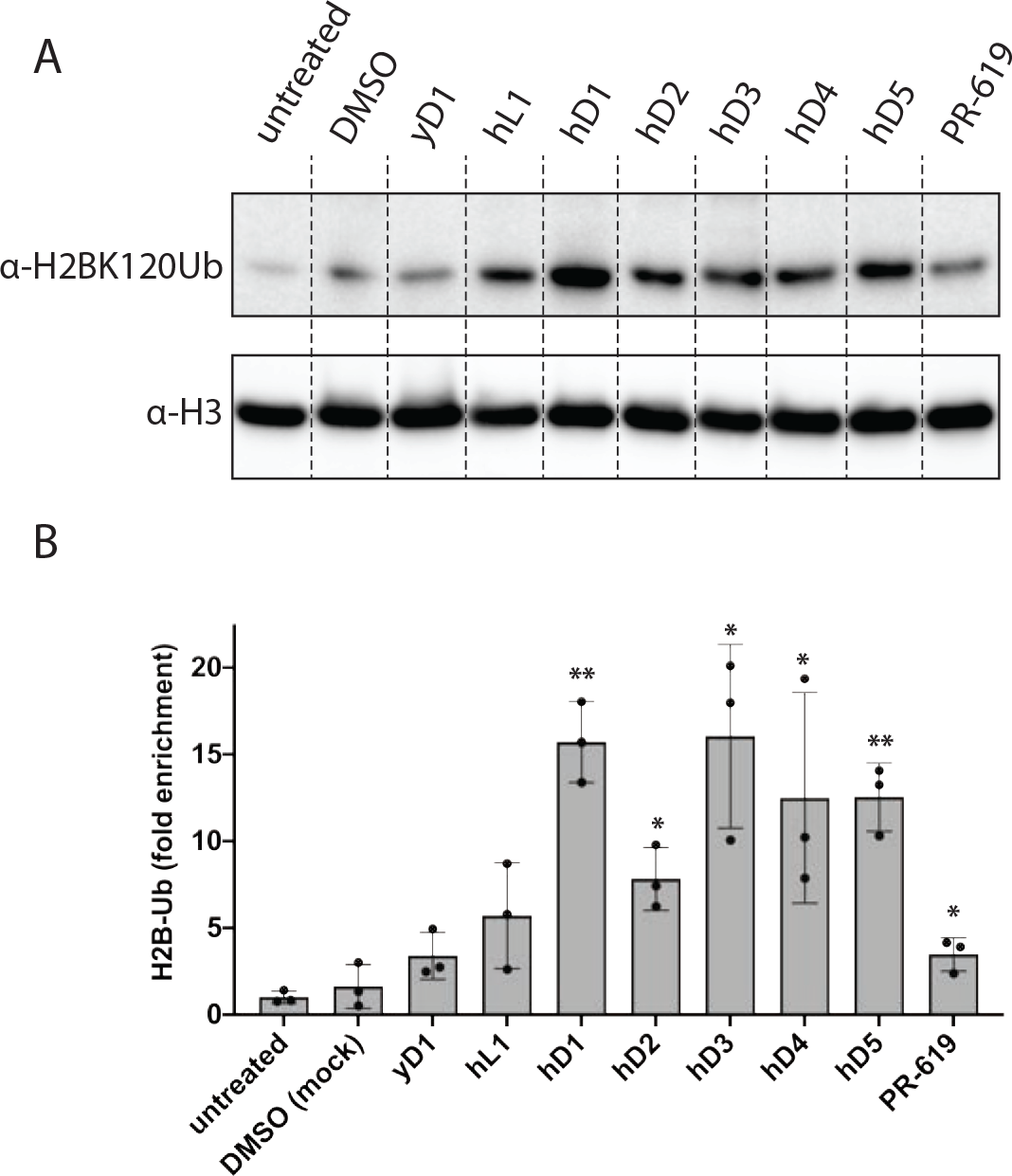
Cyclic peptide inhibitors of hDUBm increase cellular H2B-Ub levels. **A)** HEK293T cells incubated with 5 µM of each cyclic peptide for 2 hours. Histone proteins were extracted, and samples analyzed by western blot with anti-H2BK120ub antibody. Bottom: anti-histone H3 loading control. **B)** Quantitation of western replicate results indicates significant enrichment of H2B-Ub when normalized to H3 abundance. Bars labeled with two asterisks indicate p < 0.01 (hD1 and hD5), and a single asterisk indicates p < 0.05 (hD2, hD3, hD4, and PR-619) when each measurement is compared with untreated cells.

## Discussion

Specifically targeting one of the ∼80 human DUBs with inhibitors is difficult because of the high degree of structural similarity within each superfamily. Identifying specific inhibitors of the USP family is particularly challenging because there are 55 USP DUBs that share a structurally conserved catalytic domain and active site (Clague et al., 2013). Using in vitro selection with a library of cyclic peptides, we identified novel and highly specific inhibitors of USP22, a subunit of the heterotetrameric SAGA deubiquitinating module, whose overexpression is part of an 11-gene “death by cancer” signature (Glinsky et al., 2005). The most potent inhibitor, hD1, inhibits USP22 in vitro with a *K*i of 180 nM (Figure 3A). Importantly, hD1 is remarkably specific for USP22 and has little to no effect on the activity of 44 other human DUBs, with the notable exception of USP27x (Figure 5). The ability of the peptide inhibitors to increase ubiquitination in cells (Figure 6) indicates that the inhibitor is able to cross the outer cell membrane and enter the nucleus. To our knowledge, the cyclic peptides discussed in this work are the first inhibitors of the human DUB module, potentially offering opportunities for novel therapies and new insights into the roles of USP22 and H2B-Ub signaling in cancer and differentiation.

The selectivity of hD1 among DUB complexes specific for histone H2B-K120Ub is particularly striking. In addition to the SAGA DUB module, which contains USP22 in complex with ENY2, ATXN7L3 and ATXN7, two other DUB complexes have been identified that contain two of the three non-catalytic SAGA subunits and also deubiquitinate histone H2B-K120Ub (Atanassov et al., 2016). Both USP27x and USP51 form heterotrimeric complexes with ENY2 and ATXN7L3 (Atanassov et al., 2016), but lack the ATXN7 subunit that anchors the SAGA DUB module to the remainder of the SAGA complex (Herbst et al., 2021). Since hD1 inhibits USP27x whether or not it is in complex with the other SAGA subunits, the inhibitor likely binds directly to the catalytic USP domain. Out of the 55 human USP family DUBS, USP27x is most similar to USP22, with 82% sequence identify. USP51, which is only modestly inhibited by hD1 at very high concentrations, is 70% identical in sequence to USP22. The other human DUBs, including the USP DUBs tested here, are less than 40% identical in sequence to USP22. It is likely that some of the contacts formed by hD1 with the USP domain involve residues that differ among the three enzymes, with the remaining contacts involving conserved residues. Since hD1 was only active on DUBs closely related to USP22, it is likely that hD1 does not inhibit the other human DUBs that were not tested in this study.

The inhibitors tested here likely act by disrupting or occluding the active site, or by interfering with ubiquitin binding. Structural studies of the yeast DUB module showed that the USP domain contacts the ubiquitin moiety only, while the Sgf11 subunit (homologue of ATXN7L3) contacts the nucleosome acidic patch via its arginine-rich zinc finger (Kohler et al., 2010, Samara et al., 2010, Morgan et al., 2016). Substitutions of the contacting Sgf11 zinc finger arginine residues therefore abrogate DUB module activity on nucleosomes but not on Ub-AMC or diubiquitin (Morgan et al., 2016). Since the cyclic peptides we identified inhibit cleavage of both Ub-AMC and nucleosome substrates (Figure 2), the inhibitors most likely act by interfering with ubiquitin binding or catalysis. Full catalytic activity of the DUB module depends upon incorporation of the catalytic domain into the heterotetrameric complex, so an inhibitor could also potentially act by disrupting the quaternary organization of the four subunits, thereby abrogating USP22 activity (Rodriguez-Navarro et al., 2004, Ingvarsdottir et al., 2005). However, SAXS studies of the human DUB module (Figure 4) in the presence of inhibitory peptide showed no evidence of a gross conformational change of the type observed for the yeast DUB module upon deletion of the Sgf11 zinc finger (Samara et al., 2012).

Structural information will be needed to determine the binding location and mechanism of action of the DUB peptide inhibitors, as well as highlight which macrocycle residues are most important for binding to USP22. A majority of the peptides that show inhibitory activity (Figure 2A-F) contain the sequence, SYRG (hD1, hD2, hD4) or SWRG (hD5) (Figure 1); however hD3, which contains only SY, is still a potent inhibitor (Figure 2C). The macrocycle hL1, which contains an L-amino acid, rather than D-, at the N-terminal cyclization residue is unrelated in sequence to the others yet is also a potent inhibitor (Figure 2F).

Our cell-based experiments demonstrated the effectiveness of each macrocycle in enriching H2B-Ub in HEK293T cells (Figure 6). Interestingly, the effect in cells was not strictly correlated with the relative potency of each inhibitor in vitro; the weakest inhibitor of the peptides tested, hD4, enriched H2B-Ub in cells to a similar extent as the most potent inhibitor, hD1 (Figures 2 and 6A). A possible explanation is that each peptide enters cells with a different efficiency, although further study will be needed to assess peptide uptake in cells and in tissues. The mechanism by which the macrocyclic peptides described in this work enter the cells, and whether they permeate cells passively or actively, is as yet unknown. We have previously reported similar cases where the thioether-macrocyclic peptides did enter cells and actively inhibit enzyme or protein functions (Kawamura et al., 2017, Nawatha et al., 2019, Rogers et al., 2021).

Most importantly, because of the potent activity of our peptide (Ki < 200 nM) and remarkable specificity for the target, even if only a small fraction of peptide added to media could permeate to cells, they might exhibit expected cellular activity. Nevertheless, it is encouraging that treatment with macrocyclic peptide clearly enriched H2B-Ub in cells, suggesting that the inhibitor is active in cells and is membrane permeable. These results provide strong incentive to further engineer the macrocycles to improve inhibitory activity against USP22, as well as cell permeability. We note that, while the effect on cellular H2B-Ub levels observed most likely stem from inhibition of USP22, we cannot rule out a contribution from inhibiting USP27x, whose IC50 for hD1 is about 25-fold higher than for USP22. It is yet unclear the mechanism of how the macrocyclic peptides described in this work enter the cells, i.e. whether they permeate cells passively or actively, at this point.

In addition to its role in maintaining H2B-Ub levels, USP22 deubiquitinates several other proteins, including TRF1 (Atanassov et al., 2009), FBP1 (Atanassov and Dent, 2011), and SIRT1 (Lin et al., 2012), complicating the current understanding of how USP22 overexpression impacts cellular events. In mice, USP22 knockout is embryonic lethal, and hypermorphic alleles designed to reduce USP22 expression significantly altered signal transduction for agonists of several receptor tyrosine kinases, resulting in a phenotype that includes developmental aberrations in the placenta and embryo (Koutelou et al., 2019). Misregulated expression of developmental genes driving differentiated cells toward pluripotency can result in cancer progression. However the interplay between the DUB module’s roles in chromatin maintenance, transcription factor regulation, and the gene-specific consequences of those combined effects makes it difficult to attribute biological outcomes to any particular USP22 deubiquitination target (Wang and Dent, 2014). The inhibitors we have identified provide new tools to probe the role of USP22 in targeting all its cellular substrates.

## ACKNOWLEDGEMENTS

We thank Xiangbin Zhang for assistance in expressing and purifying the human DUB module and the USP27x complex. This work was supported by National Institute of General Medical Sciences grant GM130393 (C.W.) and by Grant-in-Aid for Specially Promote Research JP20H05618 (H.S.) from the Japan Society for the Promotion of Science.

## METHODS

### Protein expression and purification

For library screening, Ubp8 and USP22 constructs with N-terminal 6xHis were cloned into expression vectors, extending the N-terminus with an Avi-tag (GLNDIFEAQKIEWHE), an amino acid sequence that can be specifically biotinylated by the enzyme BirA during expression or in a reconstituted biochemical reaction. The yeast DUB module was expressed and purified as previously described (Samara et al., 2010). Briefly, bacterial cells (BL21-Rosetta2-pLysS) were transformed with three separate plasmids bearing the following coding sequences: Ubp8 (pET32a), Sgf11 and Sgf73 1- 105 (CDFDuet), and Sus1 (pRSF). Cells were harvested, resuspended in lysis buffer (50 mM HEPES pH 7.8, 500 mM NaCl, 20 mM imidazole, 50 µM ZnCl2, 5% glycerol, 15 mM betamercaptoethanol, and 1 mM PMSF) and lysed using a microfludizer (Microfluidics, Inc.) followed by passage over a HisTrap column (GE Healthcare) and elution with a 20-400 mM imidazole gradient in lysis buffer. Peak fractions were collected and dialyzed into IEX buffer (20 mM HEPES pH 7.6, 50 mM NaCl, 20 µM ZnCl2, 15 mM betamercaptoethanol, and 5% glycerol), followed by further purification with a Q-SP column (GE Healthcare) using a 0.05-1 M salt gradient in the same buffer. Fractions containing the yeast DUB module were then pooled, concentrated, and injected onto a Superdex 200 10/300 column (GE Healthcare) equilibrated with storage buffer (20 mM HEPES pH 7.6, 150 mM NaCl, 20 µM ZnCl2, 5 mM DTT, and 5% glycerol). Peak fractions were pooled, concentrate/ed and flash-frozen in aliquots.

The human DUBm was expressed using a baculovirus plasmid construct containing all four subunits, USP22, ATXN7(3-151), ATXN7L3, and ENY2, using the Bac2Bac system (Thermo), which we used to transfect Sf9 insect cells in order to amplify the resultant baculovirus. High Five cells were infected with the hDUBm baculovirus for two days, harvested, lysed, and purified in the same buffers used for yDUBm. After purification, recombinant BirA was added to the purified complexes in the presence of 50 mM biotin to produce the biotinylated complex, which was suitable for immobilization and mRNA display screening.

All biochemical and biophysical assays used yDUBm or hDUBm constructs lacking the Avi tag, which were purified essentially as described for their Avi-tagged counterparts, with the exception that the 6xHis affinity tag was cleaved from Ubp8 and USP22 by addition of TEV protease following HisTrap purification and repassage of the ensuing product over the HisTrap column to remove the cleaved tag and protease.

### Screening of DUBm binding macrocyclic peptides with the RaPID system

In vitro selections of DUBm binding macrocyclic peptides using RaPID system was performed as previously reported (Ito et al., 2015) with slight modification. Briefly, the initial random mRNA library was transcribed and ligated to a <u>puromycin</u> linker primer via T4 ligase for 30 min at 25°C and extracted with phenol/chloroform and ethanol precipitated. A 150 μL translation reaction using the methionine-deficient FIT system (Goto et al., 2011) and a 50 μM concentration of ClAc-L-Tyr-tRNA^fMet^CAU and ClAc-D- Tyr-tRNA^fMet^CAU were used to convert the mRNA library into a library of peptide-mRNA fusions. The translation was performed at 37°C for 30 min followed by a 25°C step for 12 min to enhance the formation of the peptide-mRNA fusions. Thirty microliters of 100 mM EDTA was added to dissociate ribosomes and the peptide-mRNA fusions were incubated at 37°C for 30 min to allow the <u>thioether cyclization</u> to approach completion.

The fused peptide–mRNA was subsequently reverse transcribed using MMLV RT RNase H- (Promega) for 1 h at 42 °C and 0.5 μL aliquot of the peptide-mRNA fusions was taken from the mixture and saved for the determination of the total amount of inputted mRNA. The peptide–mRNA fusions were then incubated with human and yeast DUBm-immobilized on Dynabeads M-280 streptavidin (Invitrogen) for 30 min at 37 °C. The resultant complementary DNAs were eluted by mixing with 1 × PCR reaction buffer and heating at 95 °C for 5 min, followed by immediate separation of the supernatant from the beads. A small fraction of the cDNA and input were allocated to real-time PCR quantification using a LightCycler 2.0 (Roche); the remainder was amplified by PCR. The resulting duplex DNAs were purified by phenol–chloroform extraction and ethanol precipitation, and transcribed into mRNAs for the next round of selection. From the second round of selection, the translation was performed at 5 μL scale, and six times of pre-clear steps were added as negative selection preceding the positive selection steps using 1 μL each of untreated and biotin bound Dynabeads. Finally, the observed enrichments appearing at fifth and sixth round were subjected to further DNA deep sequencing using the MiSeq sequencing system (Ilumina).

### Chemical synthesis of peptides

Macrocyclic peptides were synthesized by standard Fmoc solid-phase peptide synthesis (SPPS) using a Syro Wave automated peptide synthesizer (Biotage). The resulting peptide–resin (25 μmol scale) was treated with a solution of 92.5% trifluoroacetic acid (TFA), 2.5% water, 2.5% triisopropylsilane and 2.5% ethanedithiol, to yield the free linear *N*-ClAc-peptide. Following diethyl ether precipitation, the pellet was dissolved in 10 ml triethylamine containing DMSO and incubated for 1 h at 25 °C, to yield the corresponding macrocycle. The peptide suspensions were then acidified by addition of TFA to quench the macrocyclization reaction. The macrocycle was purified by RP-HPLC, using a Prominence HPLC system (Shimadzu) under linear gradient conditions. Mobile phase A (comprising water with 0.1% TFA) was mixed with mobile phase B (0.1% TFA in acetonitrile). Purified peptides were lyophilized *in vacuo* and molecular mass was confirmed by MALDI MS, using an AutoFlex II instrument (Bruker Daltonics).

### Ub-AMC deubiquitination assay

A working stock mixture of either yDUBm or hDUBm was prepared in the presence of either DMSO (mock) or a dilution series of the cyclic peptide. Before making the stock mix, the enzyme complex was diluted in reaction buffer (50 mM HEPES, pH 7.5, 150 mM NaCl, 1 µM ZnCl2, and 5 mM DTT), and each cyclic peptide was serially diluted with DMSO from 1.6 mM in 2-fold increments to make 15x stocks of each desired final peptide concentration (so as to keep DMSO concentration constant in all reactions), then diluted in buffer before mixing with enzyme. Each enzyme:peptide mixture was incubated for 30 min at 30°C in the case of yDUBm, and 37°C in the case of hDUBm, with each enzyme complex at 1.033x the desired concentration (207 nM). We added 29 µM of the enzyme:peptide mixture to 1 mL of 30 µM Ub-AMC (7-amino-4- methylcoumarin) (Boston Biochem), for a final concentration of 200 nM DUBm, the indicated cyclic peptide concentration, and 1 µM Ub-AMC. DUB activity was measured by observing the increase in AMC fluorescence in a plate reader, and the initial rate of fluorescence increase was calculated and plotted as a function of inhibitor concentration (Figure 2A-F). The *Ki* values were determined using the Morrison approximation of enzyme inhibition equilibria (Kuzmic et al., 2000) within Prism (Graphpad).

### Deubiquitination of nucleosomal H2B

Recombinant human mononucleosomes containing histone H2B ubiquitinated at K120 (H2B-Ub) were obtained from Epicypher (dNuc, 16-0370) and diluted to 2 µM with reaction buffer. Both hDUBm and inhibitor peptides were diluted from concentrated stocks with reaction buffer (10 mM HEPES pH 7.5, 150 mM NaCl, 1 µM ZnCl2, and 0.2 mM TCEP) to a final concentration of 400 nM DUB module and 2 µM cyclic peptide and incubated at 37°C for 20 minutes. Reactions were initiated by combining equal volumes of the enzyme-peptide mixture and nucleosomes, resulting in final concentrations of 200 nM hDUBm, 1 µM cyclic peptide, and 1 µM H2B-Ub nucleosome. Time points of the reaction were quenched with SDS sample buffer, separated by SDS-PAGE (Invitrogen, Bolt gels), and stained with Sypro Ruby (Life Technologies). Gels were imaged and quantitated (BioRad product info), using the H3 band as an independent load control to normalize the band intensities of H2B-Ub. Experiments were done in triplicate and analyzed using Prism (GraphPad).

### Small-angle x-ray scattering

Small angle x-ray scattering data (SAXS) were collected on a Rigaku BioSAXS 2000 instrument mounted on a Rigaku FRE-Super Bright rotating anode x-ray generator. The yeast and human DUBm and DUBm were diluted to concentrations of 1 mg/mL (6 µM) in reaction buffer (see H2B-Ub DUB assay) and scattering data recorded with a Pilatus- 6M detector. To record SAXS data from each DUBm in the presence of cyclic peptide, both peptide and DUBm were incubated at equimolar concentration (6 µM). Data were analyzed using the Primus suite of SAXS analysis software to obtain p(r) curves. These data were then processed further using the DENSS algorithm to produce electron density maps. Maps were then further analyzed using UCSF Chimera. (Add how you calculated theoretical curves and maps from the structure).

### DUB specificity assays

Ubiquitin-rhodamine(110)-glycine cleavage rates were measured for all DUBs available in the DUB*profiler*^TM^ panel with two independent replicates (44 in total; Ubiquigent).

Enzyme activity was measured in the presence of a constant concentration of 1 µM hD1. USP22 was not included in the panel, and is not represented in Fig 5A. Activity of purified DUB complexes targeting H2B that coopt ATXN7L3 and ENY2 was tested in the presence of a range of concentrations of hD1. Initial cleavage rates of each complex were measured on a fixed concentration of 5 µM Ub-AMC.

### Cell-based Assays

HEK293T cells were purchased from ATCC (CRL-3216) and cultures were expanded for at least 10 passages in Dulbecco’s Modified Eagle Medium (DMEM) supplemented with 10% fetal bovine serum (FBS) in T75 flasks incubated in a 37°C and 5% CO2 cabinet before any manipulation. Once cells were mature, HEK293T cells were grown to 80% confluence in 6-well plates, the media were aspirated, and media with reduced FBS (DMEM, 2% FBS) and 5 µM cyclic peptide were introduced. After two hours, the media were aspirated, and the cells were collected by scraping (∼1x10^6^ cells per well). The cells were then washed in 1x PBS twice followed by histone extraction (Active Motif). The extracted histones were then western blotted for H2B-Ub levels (Cell Signaling, 5546S), using H3 western blotting (Abcam, ab1791) to normalize loading.

## SUPPLEMENTAL FIGURES

**Supplemental Figure S1.**
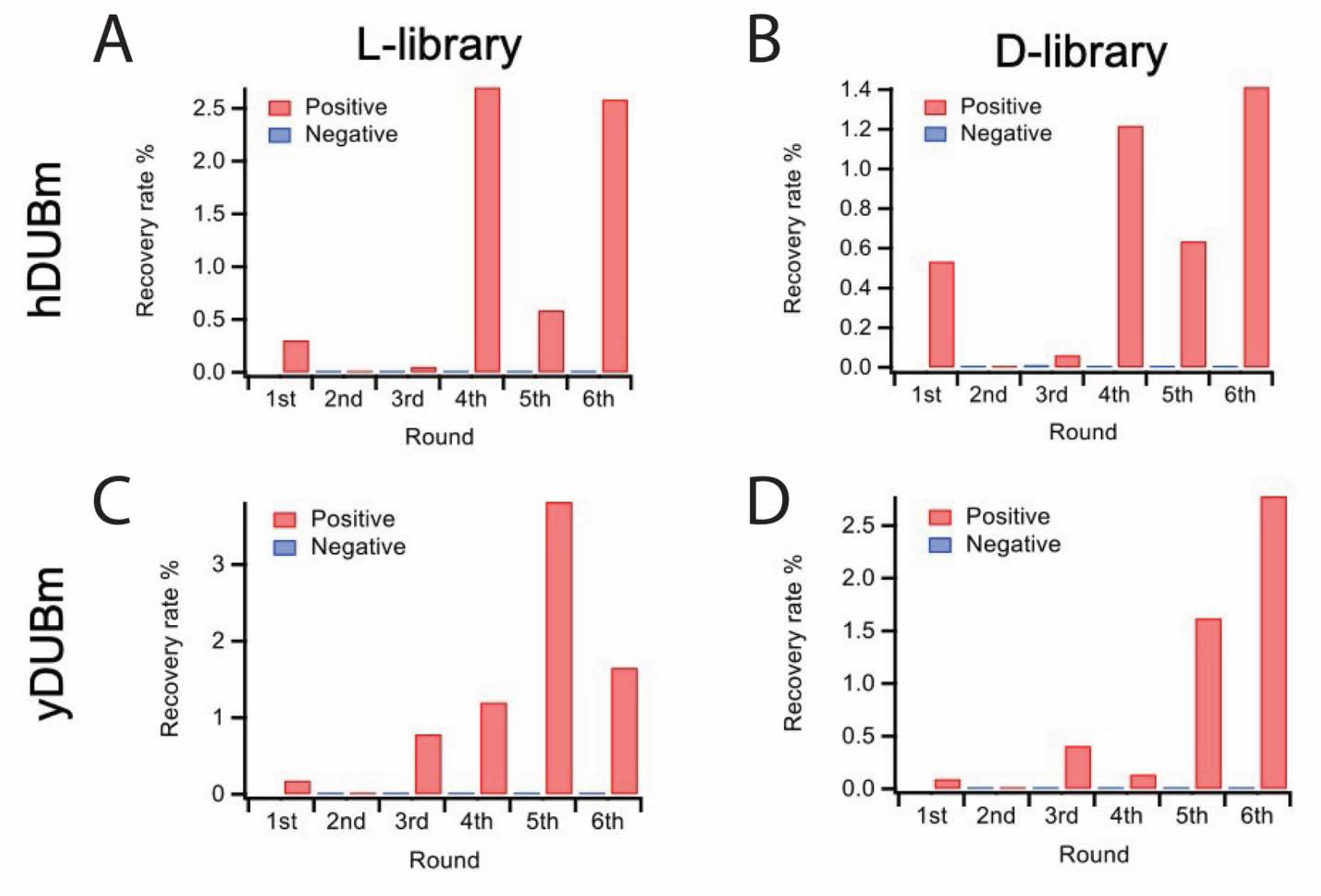
, related to Figure 1. Progress of selection for hDUBm from ^L^Y-library **A)** and ^D^Y-library **B)**, and for yDUBm from ^L^Y-library **C)** and ^D^Y-library **D)**. Red and blue bars represent the recovery rate of the cDNAs eluted from peptide-mRNA complex binding to DUBm-immobilized magnetic beads and biotin bound magnetic beads, respectively.

**Supplemental Figure S2.**
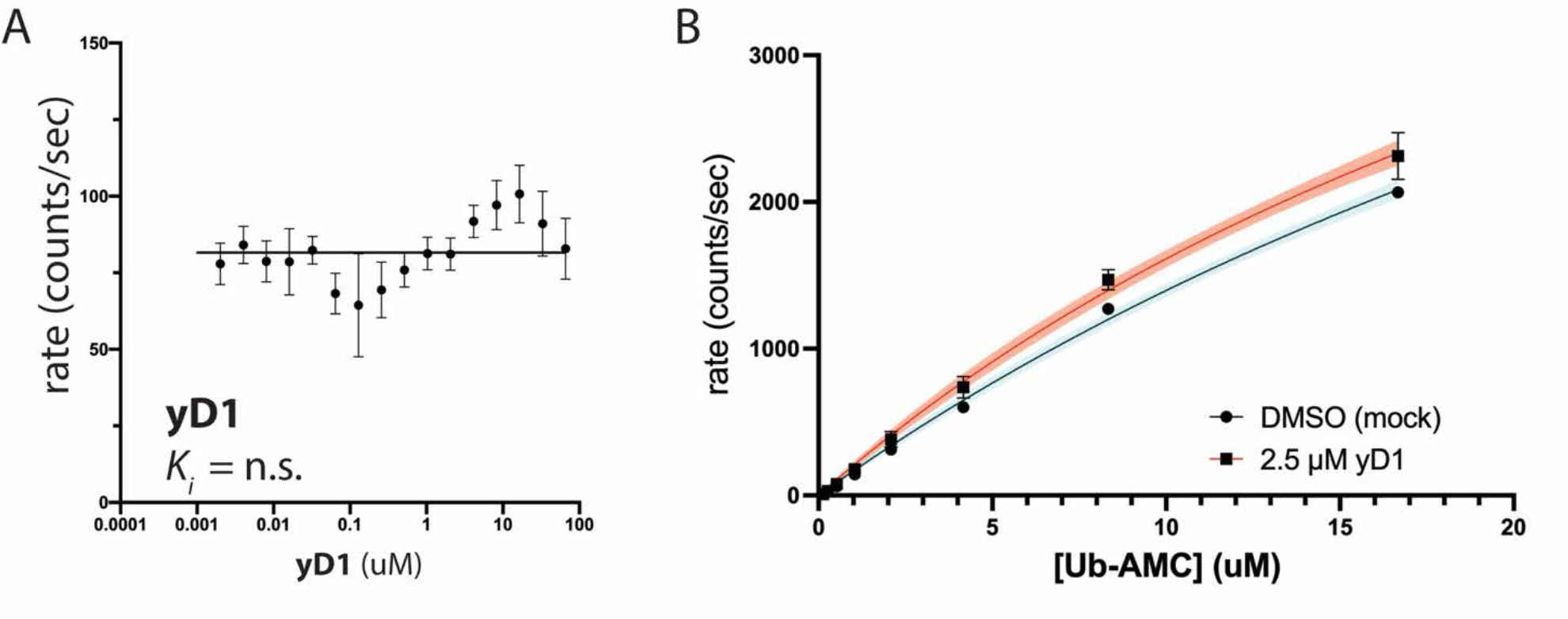
Related to Figure 2. yD1 does not inhibit the human or yeast DUB modules. A) As in Figure 2, hDUBm enzymatic activity was assayed in the presence of a range of yD1 concentrations. yD1 was selected in RaPID screening against the yeast DUB module, and thus a negative result was expected. yD1 was further used as a negative control in which a cyclic peptide is present but inactive. B) 50 nM yeast DUB module was incubated with either DMSO (vehicle) or 2.5 µM yD1 the peptide with the tightest binding to the yeast DUB module in RaPID screening. Each enzyme mixture was then mixed with a range of Ub-AMC concentrations, and cleavage was monitored by fluorescence. Rather than inhibition, yD1 produced a slight increase in cleavage rates.

**Supplemental Figure S3.**
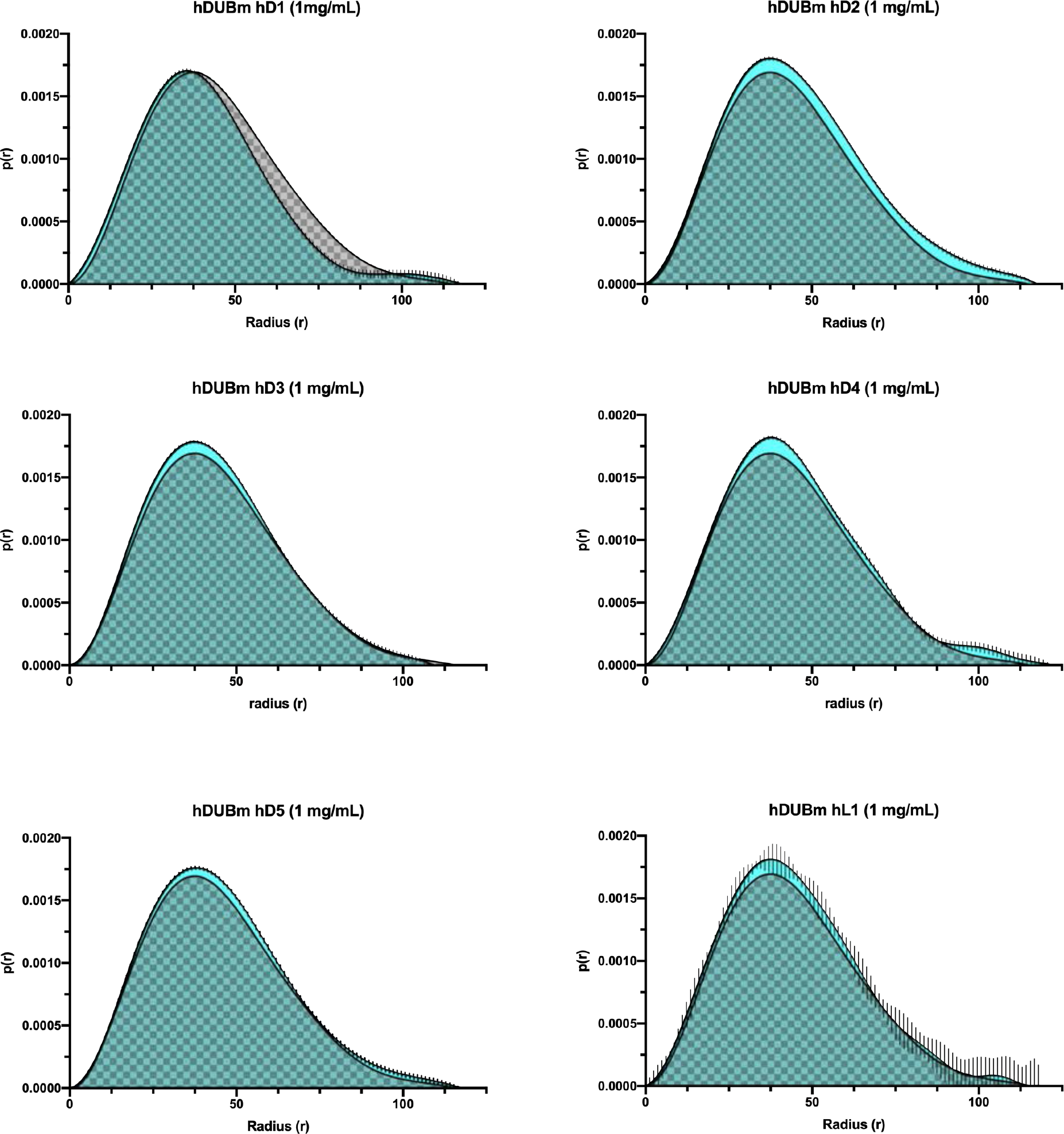
Related to Figure 3. Pairwise vector distributions [**p(r)**] calculated from SAXS data on the human DUB module collected in saturating amounts of each indicated peptide compared with data collected without peptide (checked fill). Cyan portions of the curve fill indicate differences in the p(r) distribution due to peptide binding. In the case of hD1, peptide binding apparently contracts the complex at regions indicated by the grey checkered regions.

**Supplemental Figure S4.**
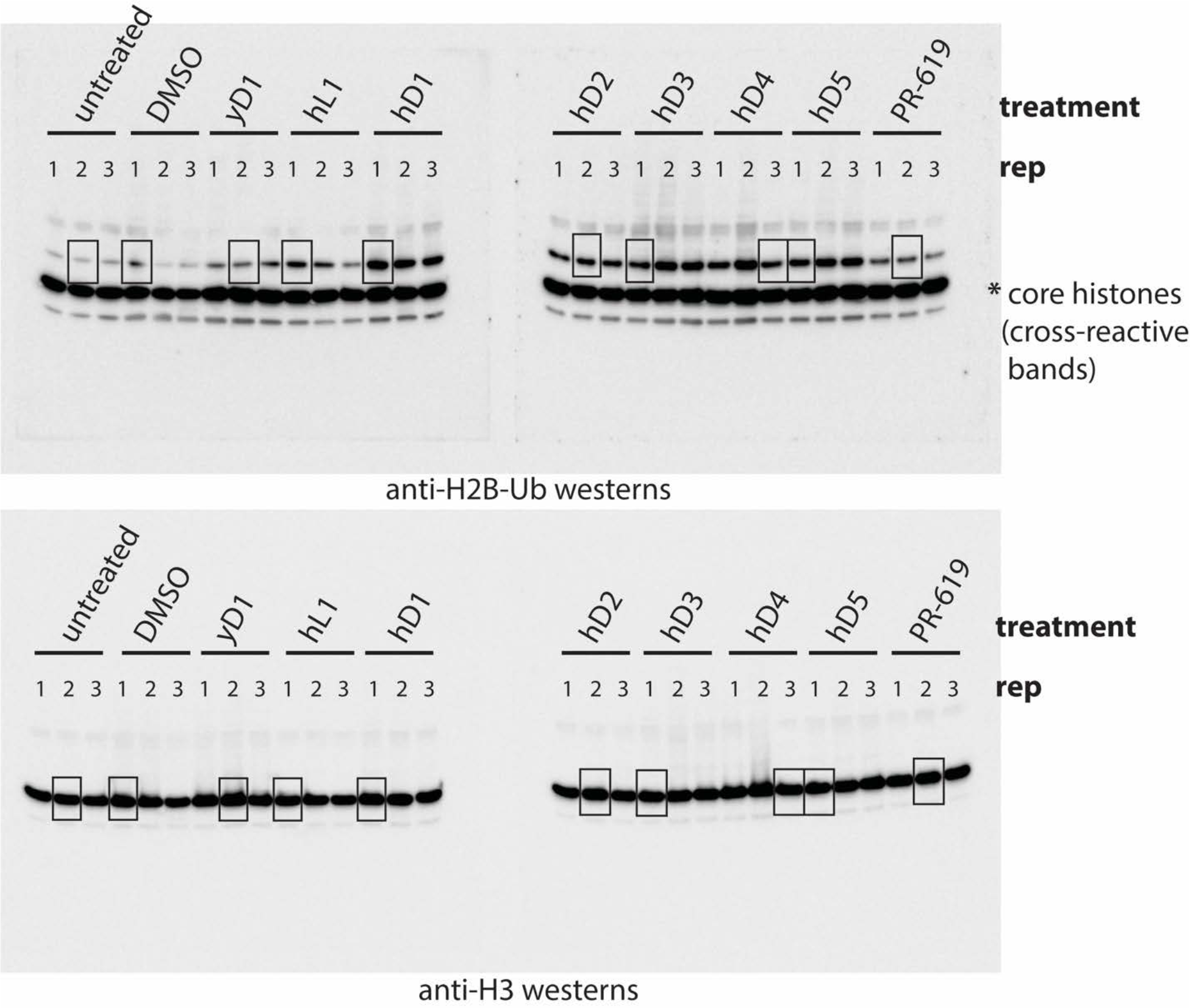
Related to Figure 4. Uncropped western blot images of anti- H2B-Ub and anti-H3 blots of identical gels normalized to H3 signals. HEK293T cells were incubated with 5 µM of each indicated peptide. Each image contains two blots from two gels imaged side-by-side with three biological replicates of the indicated treatment. Lanes cropped as representative images for Fig 4B (main text) are indicated with boxed overlays.

